# Distribution and habitat use of the Madagascar Peregrine Falcon: first estimates for area of habitat and population size

**DOI:** 10.1101/2021.08.30.458216

**Authors:** Luke J. Sutton, Lily-Arison Réné de Roland, Russell Thorstrom, Christopher J.W. McClure

**Affiliations:** The Peregrine Fund, 5668 West Flying Hawk Lane, Boise, Idaho 83709 USA; The Peregrine Fund’s Project in Madagascar, BP 4113 Antananarivo (101), Madagascar

**Keywords:** conservation planning, habitat suitability models, habitat use, population density, range maps, resource selection functions

## Abstract

Accurately demarcating species distributions has long been at the core of ecology. Yet our understanding of the factors limiting species range limits is incomplete, especially for tropical species in the Global South. Human-driven threats to the survival of many taxa are increasing, particularly habitat loss and climate change. Identifying distributional range limits of at-risk and data-limited species using Species Distribution Models (SDMs) can thus inform spatial conservation planning to mitigate these threats. The Madagascar Peregrine Falcon (*Falco peregrinus radama*) is the resident sub-species of the Peregrine Falcon complex distributed across Madagascar, Mayotte, and the Comoros Islands. Currently, there are significant knowledge gaps regarding its distribution, habitat preferences and population size. Here, we use point process regression models and ordination to identify Madagascar Peregrine Falcon environmental range limits and propose a population size estimate based on inferred habitat. From our models, the core range of the Madagascar Peregrine Falcon extends across the central upland plateau of Madagascar with a patchier range across coastal and low-elevation areas. Range-wide habitat use indicated that the Madagascar Peregrine Falcon prefers areas of high elevation and aridity, coupled with high vegetation heterogeneity and > 95 % herbaceous landcover, but generally avoids areas of > 30 % cultivated land and > 10 % mosaic forest. Based on inferred high-class habitat, we estimate this habitat area could potentially support a population size ranging between 150-300 pairs. Following International Union for Conservation of Nature Red List guidelines, we recommend this sub-species be classed as Vulnerable, due to its small population size. Despite its potentially large range, the Madagascar Peregrine has specialized habitat requirements and would benefit from targeted conservation measures based on spatial models in order to maintain viable populations.

## Introduction

On a global scale, current knowledge of species biology is heavily biased towards the Northern Hemisphere where research infrastructures have been developed over a longer time period (Meyer *et al*. 2015). In contrast, taxa inhabiting economically developing countries in tropical regions in the Southern Hemisphere often receive less attention even though these are generally the most biodiverse regions of the world (Wilson *et al*. 2016). In particular, species distributions and range limits in the Tropics are often unknown or little studied, especially for raptors (Buechley *et al*. 2019). To address distribution knowledge gaps, Species Distribution Models (SDMs) have become a widely used tool to infer species-habitat associations and identify environmental range limits (Elith & Leathwick 2009; Franklin 2009; Matthiopoulos *et al*. 2020). SDMs are statistical methods that correlate the underlying environmental conditions from known species occurrences and predict where similar environmental conditions should exist for a given species (Scott *et al*. 2002; Pearce & Boyce 2006). SDMs are particularly useful for predicting distributions for rare species inhabiting landscapes logistically and physically difficult to survey and thus estimate range size (Rhoden *et al*. 2017; Sutton & Puschendorf 2020; Sutton *et al*. 2021a,b).

The global spatial bias in mapping species distributions is most apparent for species with cosmopolitan ranges, spanning both hemispheres. The Peregrine Falcon (*Falco peregrinus*; hereafter ‘Peregrine’), is a widespread raptor with a global range, present on all continents except Antarctica (Ratcliffe 1993; White *et al*. 2013). Currently, between 18-20 geographical sub-species are recognised (White *et al*. 2013), with the distribution of many Northern Hemisphere Peregrine sub-species in Europe and North America well documented. The Madagascar Peregrine Falcon (*F. p. radama*; hereafter ‘Madagascar Peregrine’) is one of the least known Peregrine sub-species in terms of biology and distribution (White *et al*. 2013; but see Razafimanjato *et al*. 2007). This sub-species is an uncommon resident with a patchy distribution across Madagascar, Mayotte, and the Comoros Islands (Langrand 1990; Goodman *et al*. 1997; Razafimanjato *et al*. 2007). Nearly all the current literature on the Madagascar Peregrine states the scarcity of this sub-species and the logistical difficulties in locating individuals or breeding pairs (reviewed in White *et al*. 2013). Most reports state that Peregrines are most frequent in the mountainous interior of Madagascar or around rocky seacoasts and islets (Goodman *et al*. 1997; Razafimanjato *et al*. 2007).

Due to the limited transport infrastructure to survey remote areas in many regions of Madagascar, assessing distribution status for this sub-species is problematic and logistically challenging (Razafimanjato *et al*. 2007). For those species that exist in remote, hard to survey areas, SDMs are an effective method to estimate distribution using environmental layers combined with biodiversity inventory occurrence data from sources such as museums, atlases, and community science projects (Bradter *et al*. 2017; Sutton *et al*. 2021a,b). Despite issues of sampling bias in opportunistically collected inventory data (Franklin 2009; Newbold 2010), such data often cover a large sampling extent beyond that possible from systemically sampling across large scales. Thus, with improved modelling methods able to account for inherent spatial sampling biases, such as point process regression models (Renner *et al*. 2015), biodiversity inventory data can help fill knowledge gaps on distribution and habitat extent.

Defining the habitat extent and range limits of a given species is often the first step in setting conservation planning priorities (Elith & Leathwick 2009). Recently, the International Union for the Conservation of Nature (IUCN) developed a new range metric termed Area of Habitat (AOH; Brooks *et al*. 2019). Whilst this method is reproducible and widely applicable, it is unable to account for the areas where the focal species may exist but there are few or no occurrence records. Using predictive spatial models calibrated with the AOH parameters can account for these disparities, providing further detail beyond what the AOH metric can provide (Breiner *et al*. 2017; Herkt *et al*. 2017; Sutton *et al*. 2021b). Here, we use point process logistic regression and environmental ordination in an SDM framework as described by Sutton *et al*. (2021b) using Resource Selection Functions (RSFs) and Habitat Suitability Models (HSMs). Both RSFs and HSMs are conceptually the same method, under the general SDM analytical paradigm of predicting species distributions based on species-habitat associations (Boyce & McDonald 1999; Kearney 2006; Matthiopoulos *et al*. 2020). Specifically, our aims are to address three significant knowledge gaps for the Madagascar Peregrine: (**1**) provide the first detailed distribution map and area of habitat, (**2**) define habitat requirements across the current known range, and (**3**) calculate a first estimate of population size based on inferred habitat area.

## Methods

### Study Area

Madagascar, Mayotte, and the Comoros are situated in the tropical zone approximately 400 km from the south-east African mainland in the Indian Ocean. Madagascar is a continental island separated from the African mainland by the Mozambique channel, with Mayotte and the Comoros a small archipelago of volcanic islands lying 200 km off north-west Madagascar in the northern Mozambique channel (Fig. 1). Madagascar has a highly heterogeneous landscape with an extensive upland plateau running longitudinally along the centre of island, descending to a wide lowland dry forest coastal plain in the west, an arid region in the south-west, and a narrower coastal strip of humid tropical forest running along the eastern seaboard. Due to this varied landscape and topography, climate is dominated by the north-south central plateau, dictating temperature and rainfall regimes in multiple bioclimatic regions (Jolly *et al*. 2016). Both Mayotte and the Comoros have similar environmental heterogeneity, albeit at a smaller scale, with upland areas, coastal plains, and rocky coasts.

**Figure 1.**
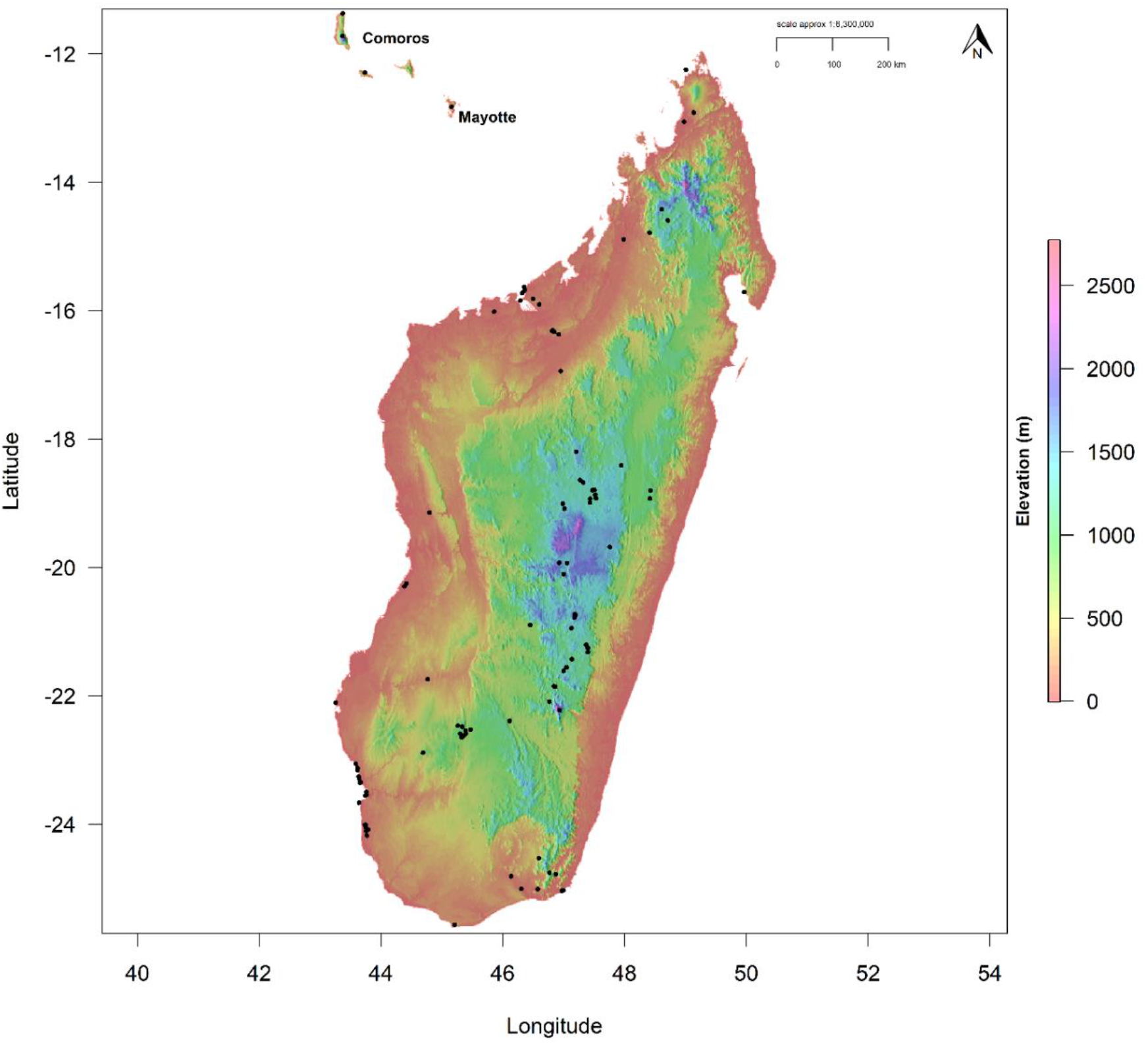
Digital Elevation Model (DEM) for Madagascar, Mayotte and the Comoros, showing altitudinal range within the current known distribution of the Madagascar Peregrine Falcon. Black points denote Peregrine occurrences.

### Occurrences

We sourced Madagascar Peregrine occurrences from the Global Raptor Impact Network (GRIN) information system (McClure *et al*. In press). For the Madagascar Peregrine, GRIN consists of occurrences from the African Raptor DataBank (Davies *et al*. 2020), eBird (Sullivan *et al*. 2009), and GBIF (2019a,b). We cleaned occurrence data by removing duplicate records, those with no georeferenced location, and any locations over water, resulting in 194 occurrences to use in the calibration models. Only occurrences recorded from 1990 onwards were used to match the timeframe of the environmental data, whilst retaining sufficient sample size for modelling (van Proosdij *et al*. 2016). We reduced sampling bias and clustering in the occurrence points, by applying a 1-km spatial filter between the raw occurrences to minimise the effects of over-sampling in highly surveyed areas (Aiello-Lammens *et al*. 2015). We selected the 1-km thinning distance to match the resolution of the environmental raster layers and to capture the high environmental heterogeneity of the study area. Removing clustered occurrence points reduces model over-fitting, minimizes omission errors and improves model predictive performance (Kramer-Schadt *et al*. 2013; Boria *et al*. 2014; Radosavljevic & Anderson 2014), and is more effective than other methods of spatial bias correction (Fourcade *et al*. 2014).

### Habitat covariates

We predicted occurrence using habitat covariates representing climate, landcover, topography and vegetation heterogeneity downloaded from the EarthEnv (www.earthenv.org) and ENVIREM (Title & Bemmels 2018) data repositories. We used a total of six continuous covariates (Table 1) at a spatial resolution of 30 arc-seconds (~1 km resolution), cropped to a delimited polygon consisting of all three range countries. We selected covariates *a prioiri* based on climate, landcover and topographic variables related empirically to Peregrine distribution (Ratcliffe 1993; White *et al*. 2013). Heterogeneity is a similarity measure for Enhanced Vegetation Index (EVI) between adjacent pixels; sourced from the Moderate Resolution Imaging Spectroradiometer (MODIS, https://modis.gsfc.nasa.gov/). We inverted the raster cell values in the original EarthEnv variable ‘Homogeneity’ (Tuanmu & Jetz 2015) to represent vegetation heterogeneity which varies between zero (minimum heterogeneity) and one (maximum heterogeneity).

**Table 1.**
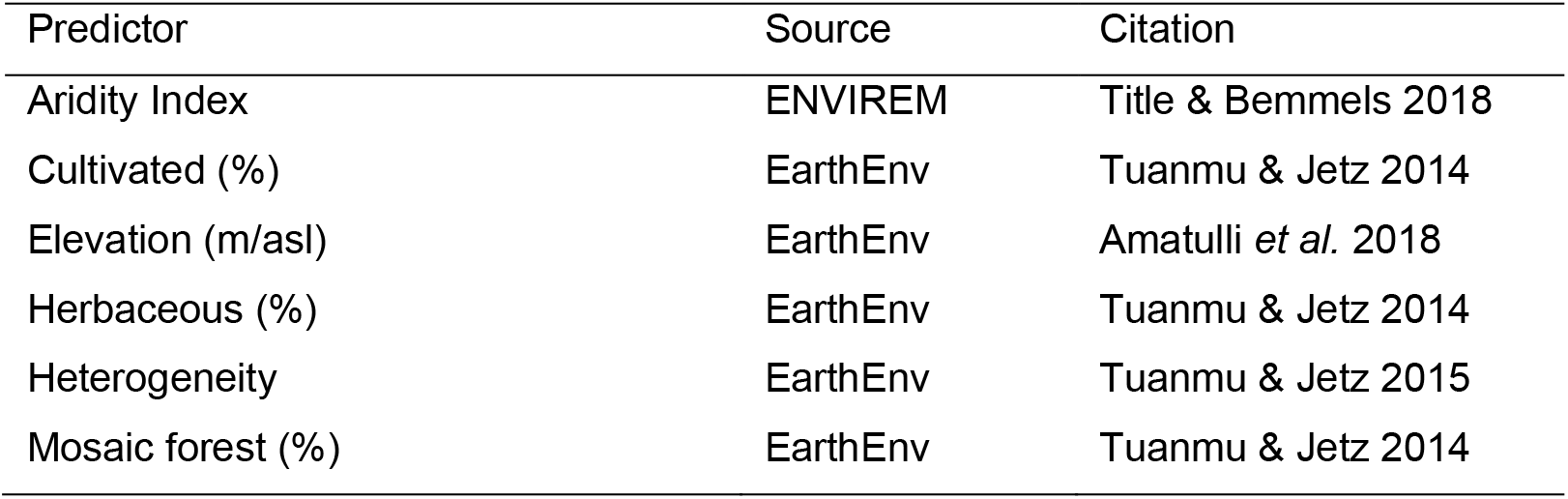
Habitat covariates used in all spatial analyses for the Madagascar Peregrine Falcon.

| Predictor | Source | Citation |
| --- | --- | --- |
| Aridity Index | ENVIREM | Title & Bemmels 2018 |
| Cultivated (%) | EarthEnv | Tuanmu & Jetz 2014 |
| Elevation (m/asl) | EarthEnv | Amatulli *et al.* 2018 |
| Herbaceous (%) | EarthEnv | Tuanmu & Jetz 2014 |
| Heterogeneity | EarthEnv | Tuanmu & Jetz 2015 |
| Mosaic forest (%) | EarthEnv | Tuanmu & Jetz 2014 |

The three measures of percentage landcover (Cultivated, Mosaic forest, Herbaceous vegetation) are consensus products integrating GlobCover (v2.2), MODIS land-cover product (v051), GLC2000 (v1.1) and DISCover (v2) from the years 1992-2006. Mosaic forest represents a mixed landcover of forest, shrubland and grassland. Herbaceous vegetation defines both open and closed grassland cover. Full details on methodology and image processing can be found in Tuanmu & Jetz (2014). Elevation was derived from a digital elevation model (DEM) product from the 250m Global Multi-Resolution Terrain Elevation Data 2010 (GMTED2010, Danielson & Gesch 2011). Aridity Index is based on the degree of water deficit below a given need (Thornthwaite 1948). The index ranges from zero (low aridity) to one (high aridity). All selected covariates showed low collinearity and thus all six were included as predictors in model calibration (Variance Inflation Factor < 2; Table S1). Lastly, we extracted the underlying environmental data for each covariate at all filtered location points to provide a summary of environmental tolerances across the Madagascar Peregrine range.

### Resource Selection Functions

We followed the spatial modelling framework described by Sutton *et al*. (2021b) and used presence points and six covariates to fit an RSF using logistic regression with a binomial error term and logit link function using a generalised linear model (GLM) in the R stats package (R Core Team 2018). The RSF followed geographical range first-order selection (Johnson 1980) using design I in a use-availability sampling protocol (Manly *et al*. 2002; Thomas & Taylor 2006). Linear and quadratic terms were fitted dependent on the scaled responses from fitting both terms on an initial model. Only linear terms were used when the quadratic term resulted in biologically unrealistic U-shaped curves, or when a linear term was sufficient to explain the scaled response.

The RSF was fitted to derive maximum likelihood estimates on model parameters significantly different from zero, with no interaction terms. Predictors were standardized with a mean of zero and standard deviation of one. Because the occurrence data correspond to a presence-only dataset, background absences representing availability were randomly sampled using 10,000 points (Barbet-Massin *et al*. 2012), and equal weights assigned to both presence and background points. To test calibration accuracy, the explained variance from each logistic model was measured using McFadden’s adjusted *R^2^* (*R^2^_adj_*, McFadden 1974). Lastly, partial response curves based on the standardized covariates of the fitted model were plotted against the scaled responses with 95 % Confidence Intervals.

We performed an Ecological Niche Factor Analysis (ENFA, Hirzel *et al*. 2002) as a second multivariate resource selection method, similar to a Principal Component Analysis (PCA), quantifying two measures of resource selection in environmental space along two axes. The first axis metric, marginality (*M*), measures the position of the species ecological niche, and its departure relative to the available environment. A value of *M* >1 indicates that the niche deviates more relative to the reference environmental background and has specific environmental preferences compared to the available environment. The second axis metric, specialization (*S*), is an indication of niche breadth size relative to the environmental background, with a value of *S* >1 indicating higher niche specialization (narrower niche breadth). A high specialization value indicates a strong reliance on the environmental conditions that mainly explain that specific dimension. ENFA was performed in the R package CENFA (Rinnan 2018), using the unfiltered occurrences (as spatial clustering is not an issue; Basille *et al*. 2008), and weighting all cells by the number of observations (Rinnan & Lawler 2019).

### Habitat Suitability Model

We fitted HSMs using penalized elastic-net logistic regression (Zou & Hastie 2005; Fithian & Hastie 2013), in the R packages glmnet (Friedman *et al*. 2010) and maxnet (Phillips *et al*. 2017). Elastic net logistic regression imposes a penalty (known as regularization) to the model shrinking the coefficients of variables that contribute the least towards zero (or exactly zero), to reduce model complexity (Gastón & García-Viñas 2011; Helmstetter *et al*. 2020). An elastic net is used to perform automatic variable selection and continuous shrinkage simultaneously (Zou & Hastie 2005; Phillips *et al*. 2017), retaining all variables that contribute less by shrinking coefficients to either exactly zero or close to zero. The complementary log-log (cloglog) transform was selected as a continuous index of environmental suitability, with 0 = low suitability and 1 = high suitability. A random sample of 10,000 background points were used as pseudo-absences recommended for regression-based modelling (Barbet-Massin *et al*. 2012) and to sufficiently sample the background calibration environment (Guevara *et al*. 2018).

Optimal-model selection was based on Akaike’s Information Criterion (Akaike 1974) corrected for small sample sizes (AIC_c_; Hurvich & Tsai 1989), to determine the most parsimonious model from two model parameters: regularization multiplier (β) and feature classes (Warren & Seifert 2011). Eighteen candidate models of varying complexity were built by comparing a range of regularization multipliers from 1 to 5 in 0.5 increments, and two feature classes (response functions: Linear, Quadratic) in all possible combinations using the ‘checkerboard2’ method of cross-validation (*k*-folds *=* 5) within the R package ENMeval (Muscarella *et al*. 2014). All models with a ΔAIC_c_ < 2 were considered as having strong support (Burnham & Anderson 2004), and the model with the lowest ΔAIC_c_ selected. Response curves and parameter estimates were used to measure variable performance within the optimal calibration model.

We used Continuous Boyce index (CBI, Hirzel *et al*. 2006) as a measure of how predictions differ from a random distribution of observed presences (Boyce *et al*. 2002). CBI is consistent with a Spearman correlation (*r_s_*) with CBI values ranging from −1 to +1, with positive values indicating predictions consistent with observed presences, values close to zero no different than a random model, and negative values indicating areas with frequent presences having low environmental suitability. CBI was calculated using 20 % test data with a moving window for threshold-independence and 101 defined bins in the R package enmSdm (Smith 2019). We tested the optimal model against random expectations using partial Receiver Operating Characteristic ratios (pROC), which estimates model performance by giving precedence to omission errors over commission errors (Peterson *et al*. 2008). Partial ROC ratios range from 0 – 2 with 1 indicating a random model. Function parameters were set with a 10 % omission error rate, and 1000 bootstrap replicates on 50 % test data to determine significant (*α* = 0.05) pROC values >1.0 in the R package ENMGadgets (Barve & Barve, 2013).

### Reclassified discrete model

To predict area of habitat, the continuous prediction was reclassified to a threshold prediction using three discrete quantile classes representing habitat suitability (Low: 0.000 - 0.378; Medium: 0.379 - 0.648; High: 0.649 - 1.000). Class-level landscape metrics were calculated on the discrete quantile classes to estimate total and core area of habitat in each class in the R package SDMTools (VanDerWal *et al*. 2014), based on the fragstats program (McGarigal *et al*. 2002). Core areas were defined as those cells with edges wholly within each habitat class, with cells containing at least one adjacent edge to another class cell considered as edge habitat. Peregrine pair densities vary widely across their global range (White *et al*. 2013) and no density estimates are currently available for the Madagascar Peregrine. Therefore, we used pair densities ranging from one pair per 500 km^2^ based on studies from Europe (Ratcliffe 1993) and one pair per 1000 km^2^ from the African mainland (White *et al*. 2013) to calculate a population size estimate from inferred habitat area. Geospatial analysis, modelling and visualisation were conducted in R (v3.5.1; R Core Team, 2018) using the dismo (Hijmans *et al*. 2017), raster (Hijmans 2017), rgdal (Bivand *et al*. 2019), rgeos (Bivand & Rundle 2019), sp (Bivand *et al*. 2013) and wesanderson (Ram & Wickham 2018) packages.

## Results

Madagascar Peregrine occurrences were located in areas of medium to high aridity, at a median elevation of 646 m/asl, but up to 2439 m/asl, with a wide inter-quantile range (Table 2). Madagascar Peregrine occurrences were associated with medium to high areas of heterogenous vegetation, but at lower percent mosaic forest (median = 24.5 %), herbaceous (median = 14 %) and cultivated (0 %) land cover but within a broad range of minimum and maximum percentages.

**Table 2.** Summary statistics from the habitat covariates underlying occurrences for the Madagascar Peregrine Falcon. IQR = Inter quartile range.

| Covariate | Median | IQR | Min. | Max. |
| --- | --- | --- | --- | --- |
| Aridity Index | 69.0 | 19.6 | 25.2 | 83.8 |
| Cultivated (%) | 0.0 | 21.0 | 0.0 | 61.0 |
| Elevation (m) | 645.5 | 1185.0 | 0.0 | 2439.0 |
| Herbaceous (%) | 14.0 | 31.3 | 0.0 | 100.0 |
| Heterogeneity (0.0-1.0) | 0.66 | 0.20 | 0.30 | 0.92 |
| Mosaic forest (%) | 24.5 | 30.3 | 0.0 | 97.0 |

### Resource Selection Functions

The GLM explained 89 % of the variability in habitat use (McFadden’s *R^2^_adj_* = 0.89). The Madagascar Peregrine was most likely to be positively associated with higher aridity and elevation and high vegetation heterogeneity (Table 3; Fig. 2). The strongest negative associations were more likely with mosaic forest and cultivated land, but with a flatter quadratic curve for herbaceous cover. In environmental space, the ENFA deviated marginally from the average background habitat available (Fig. 3), with the marginality factor lower than the average background habitat (*M* = 0.536). The Madagascar Peregrine was restricted to specific habitat relative to the background habitat with specialized habitat requirements (*S* = 1.101).

**Figure 2.**
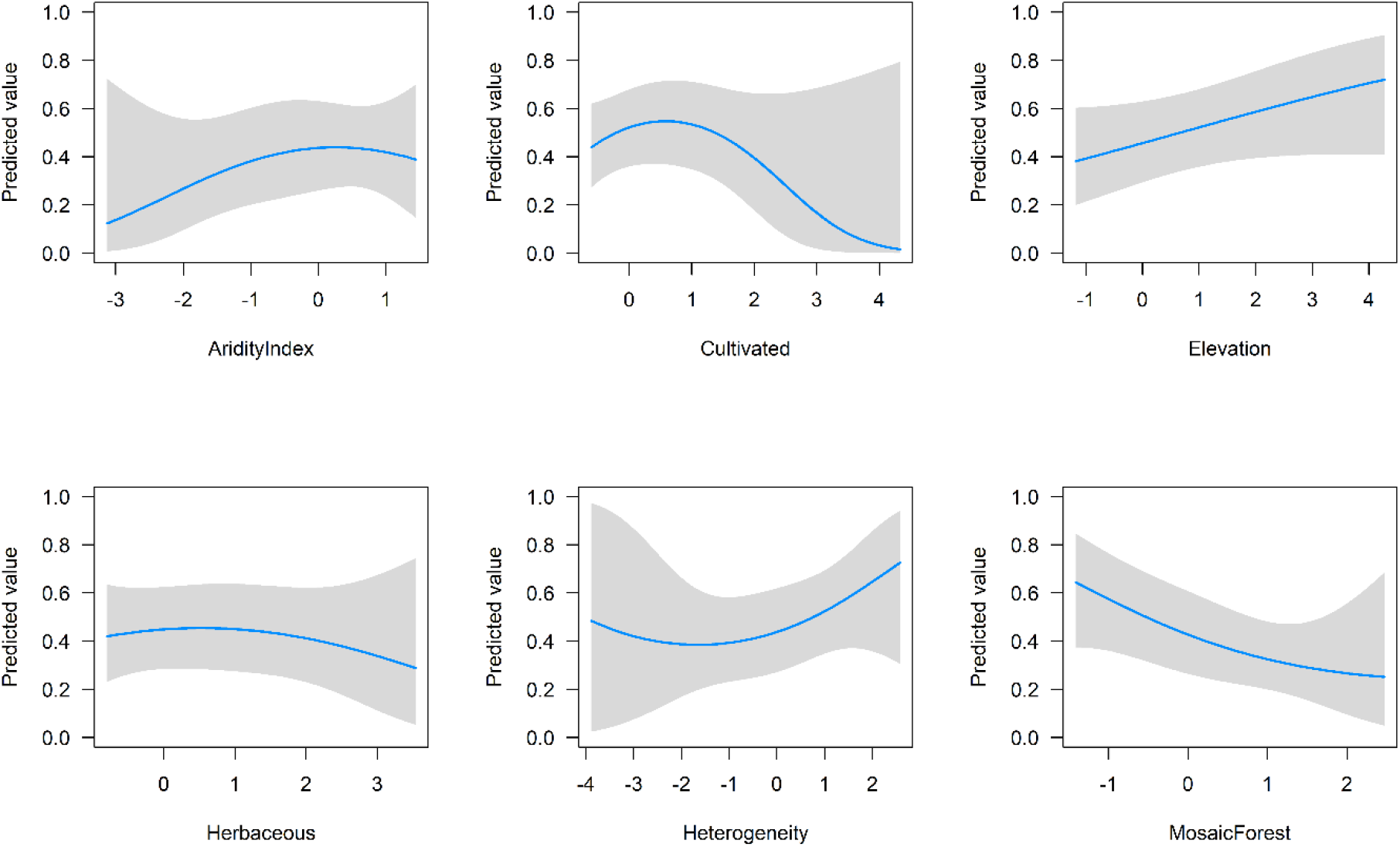
Scaled partial response curves with 95 % Confidence Intervals (grey shading) derived from maximum likelihood estimates obtained from the RSF. X-axis values are the standardized responses for each covariate (mean = 0, variance = 1).

**Figure 3.**
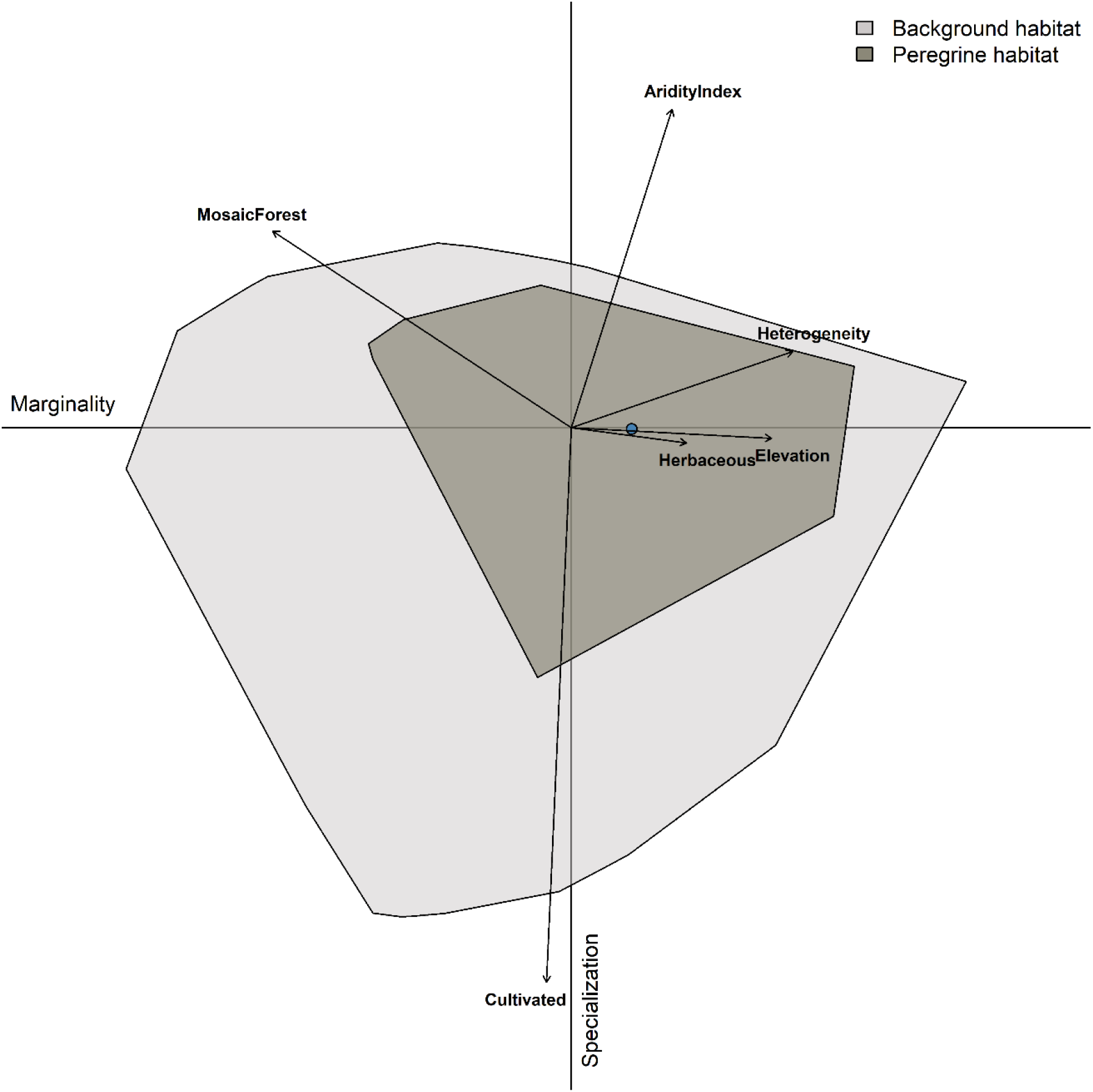
Multivariate resource selection in environmental space using Ecological Niche Factor Analysis (ENFA). Peregrine habitat (dark grey) is represented within the available background environment (light grey) shown across the marginality (x) and specialization (y) axes. Arrow length indicates the magnitude with which each variable accounts for the variance on each of the two axes. Blue point indicates niche position (median marginality) relative to the average background environment (the plot origin).

**Table 3.**
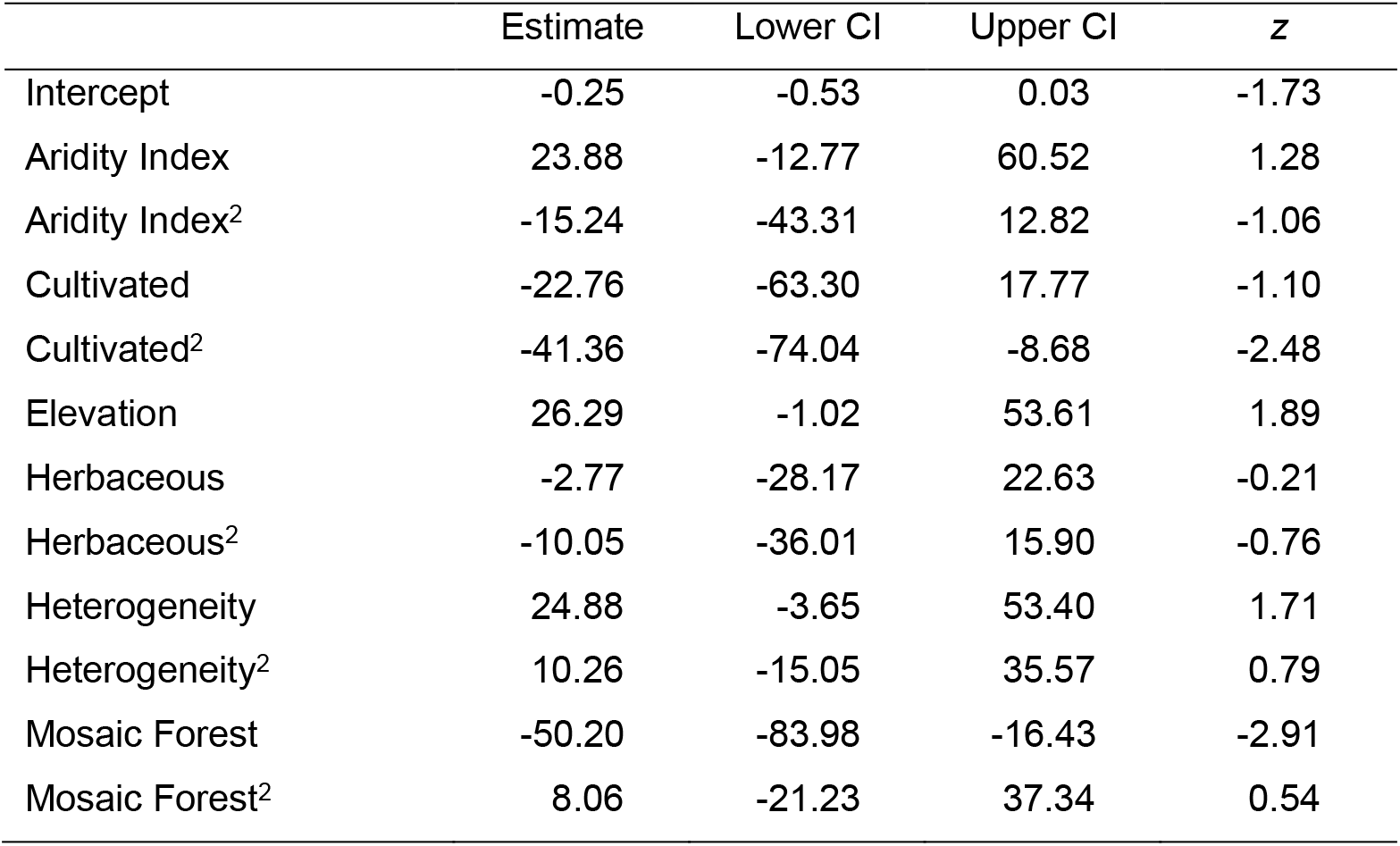
Linear and Quadratic (defined with superscript 2) terms derived from GLM maximum likelihood estimates obtained from the full model with 95 % Confidence Intervals.

|  | Estimate | Lower CI | Upper CI | z |
| --- | --- | --- | --- | --- |
| Intercept | -0.25 | -0.53 | 0.03 | -1.73 |
| Aridity Index | 23.88 | -12.77 | 60.52 | 1.28 |
| Aridity Index<sup>2</sup> | -15.24 | -43.31 | 12.82 | -1.06 |
| Cultivated | -22.76 | -63.30 | 17.77 | -1.10 |
| Cultivated<sup>2</sup> | -41.36 | -74.04 | -8.68 | -2.48 |
| Elevation | 26.29 | -1.02 | 53.61 | 1.89 |
| Herbaceous | -2.77 | -28.17 | 22.63 | -0.21 |
| Herbaceous<sup>2</sup> | -10.05 | -36.01 | 15.90 | -0.76 |
| Heterogeneity | 24.88 | -3.65 | 53.40 | 1.71 |
| Heterogeneity<sup>2</sup> | 10.26 | -15.05 | 35.57 | 0.79 |
| Mosaic Forest | -50.20 | -83.98 | -16.43 | -2.91 |
| Mosaic Forest<sup>2</sup> | 8.06 | -21.23 | 37.34 | 0.54 |

### Habitat Suitability Model

Three candidate models had an ΔAIC_c_ ≤ 2 (Table S2), and the model with the lowest ΔAIC_c_ was selected. The best-fit HSM (ΔAIC_c_ = 0.00) had linear and quadratic terms and β = 3 as model parameters, with high calibration accuracy (CBI = 0.847), and was robust against random expectations (pROC = 1.599, SD± 0.118, range: 1.141 – 1.920). The Madagascar Peregrine was most positively associated with vegetation heterogeneity, cultivated land, and highly arid areas, and negatively associated with mosaic forest (Table 4). The largest continuous area of habitat extended across the central plateau (Fig. 4). A second large area of habitat was identified across the northern Tsaratanana massif. From the HSM response curves, aridity index had peak suitability at 90 %, with highest suitability for areas of increased elevation >1500 m (Fig. 5). Habitat suitability was highest in areas of high vegetation heterogeneity > 0.9, areas with ~30 % human cultivated landcover, and < 10 % of mosaic forest cover.

**Figure 4.**
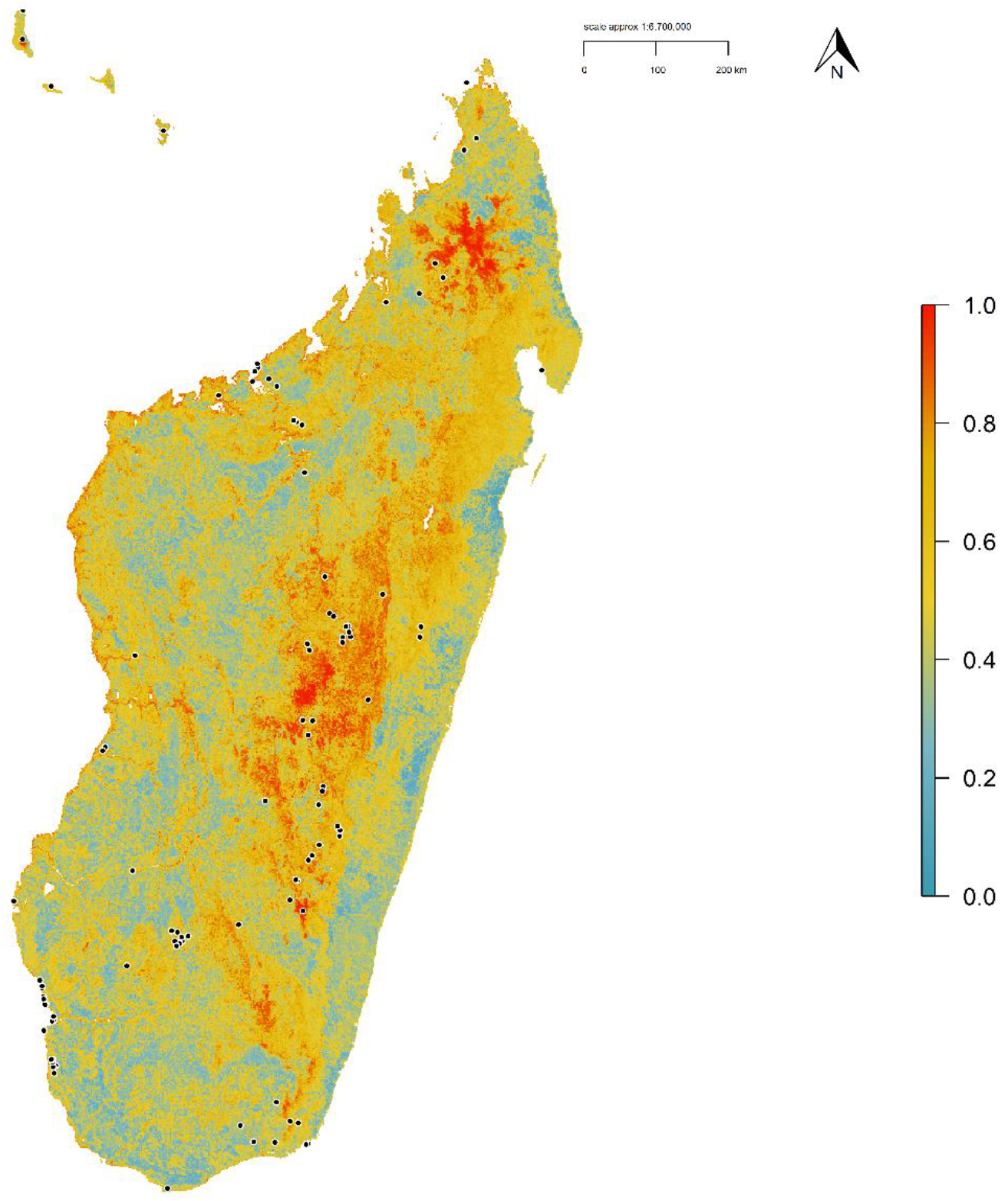
Habitat Suitability Model for the Madagascar Peregrine Falcon. Map denotes cloglog prediction with red areas having higher habitat suitability. Black points define Peregrine occurrences.

**Figure 5.**
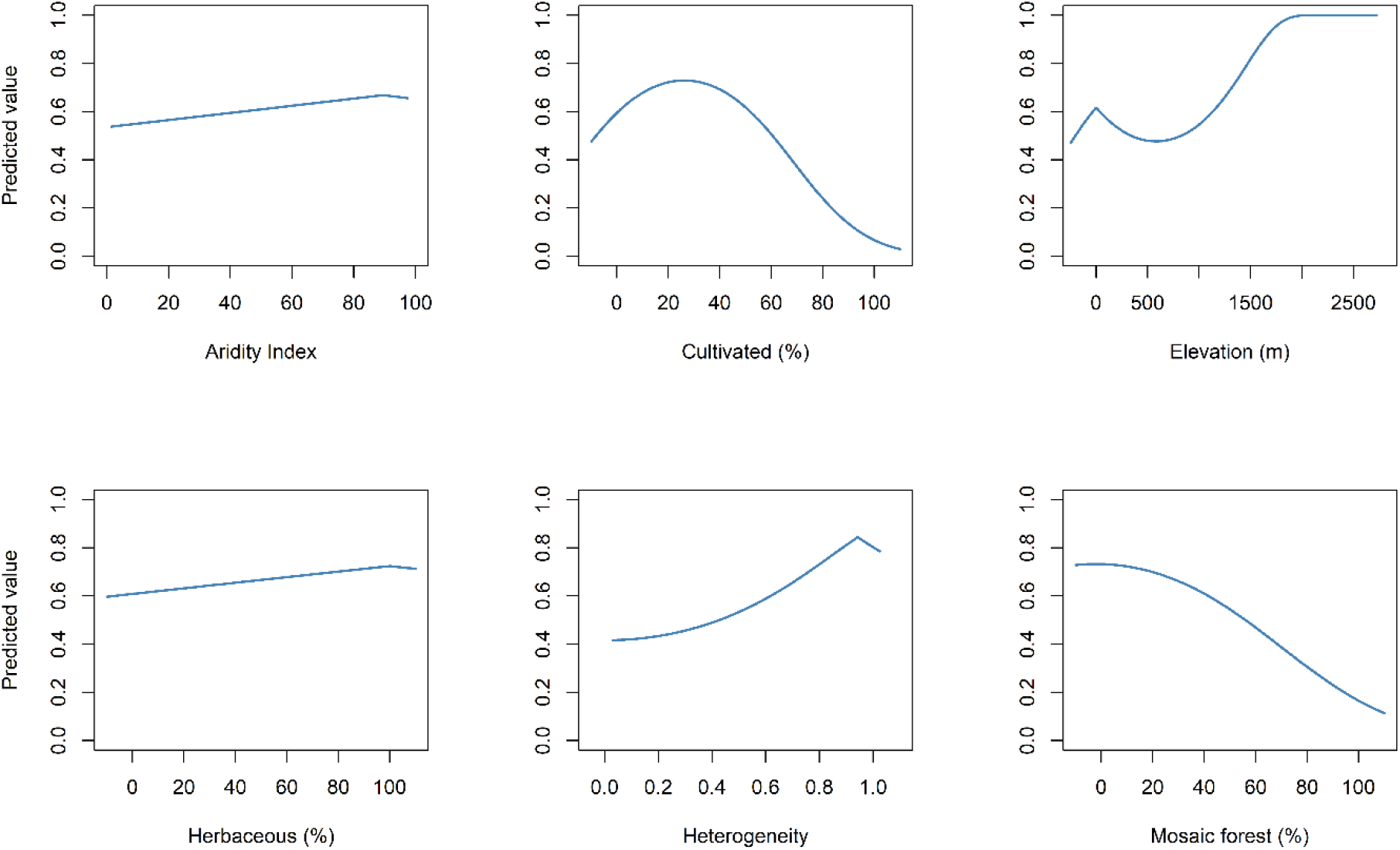
Covariate response curves from the Habitat Suitability Model for the Madagascar Peregrine Falcon.

**Table 4.** Parameter estimates derived from the HSM penalized elastic net regression beta coefficients.

| Covariate | Linear | Quadratic |
| --- | --- | --- |
| Heterogeneity | 1.875 |  |
| Cultivated | 0.018 | 0.000 |
| Mosaic forest | -0.017 |  |
| Aridity Index | 0.012 |  |
| Elevation | 0.000 | 0.000 |
| Herbaceous |  | 0.000 |

### Area of habitat and population size

The reclassified binary model calculated an area of 153,265 km^2^ in the high habitat quantile class (≥ 0.649), comprising 26 % of the total landscape area (Table 5). Core areas of high-class habitat totalled 37,466 km^2^, comprising 24.5 % of the total high habitat area. Based on the total area of high-class habitat and assuming pair densities ranging between 500-1000 km^2^ per territorial pair, we estimate the area of high-class habitat could support between 153-307 breeding pairs. From our population estimate we recommend the Madagascar Peregrine sub-species be classed as Vulnerable under IUCN Red List criterion D1 with a very small or restricted population numbering < 1000 mature individuals (IUCN 2019).

**Table 5.** Class-level landscape metrics calculated from the reclassified Habitat Suitability Model quantile threshold classes. Area values are in km^2^.

| Habitat class | Quantile threshold | Total area (%) | Core area |
| --- | --- | --- | --- |
| High | >0.649 | 153,265 (0.26) | 37,466 |
| Medium | 0.379-0.648 | 230,888 (0.39) | 12,251 |
| Low | <0.378 | 209,387 (0.35) | 43,809 |
| Total |  | 593,540 (100) | 93,526 |

**Figure 6.**
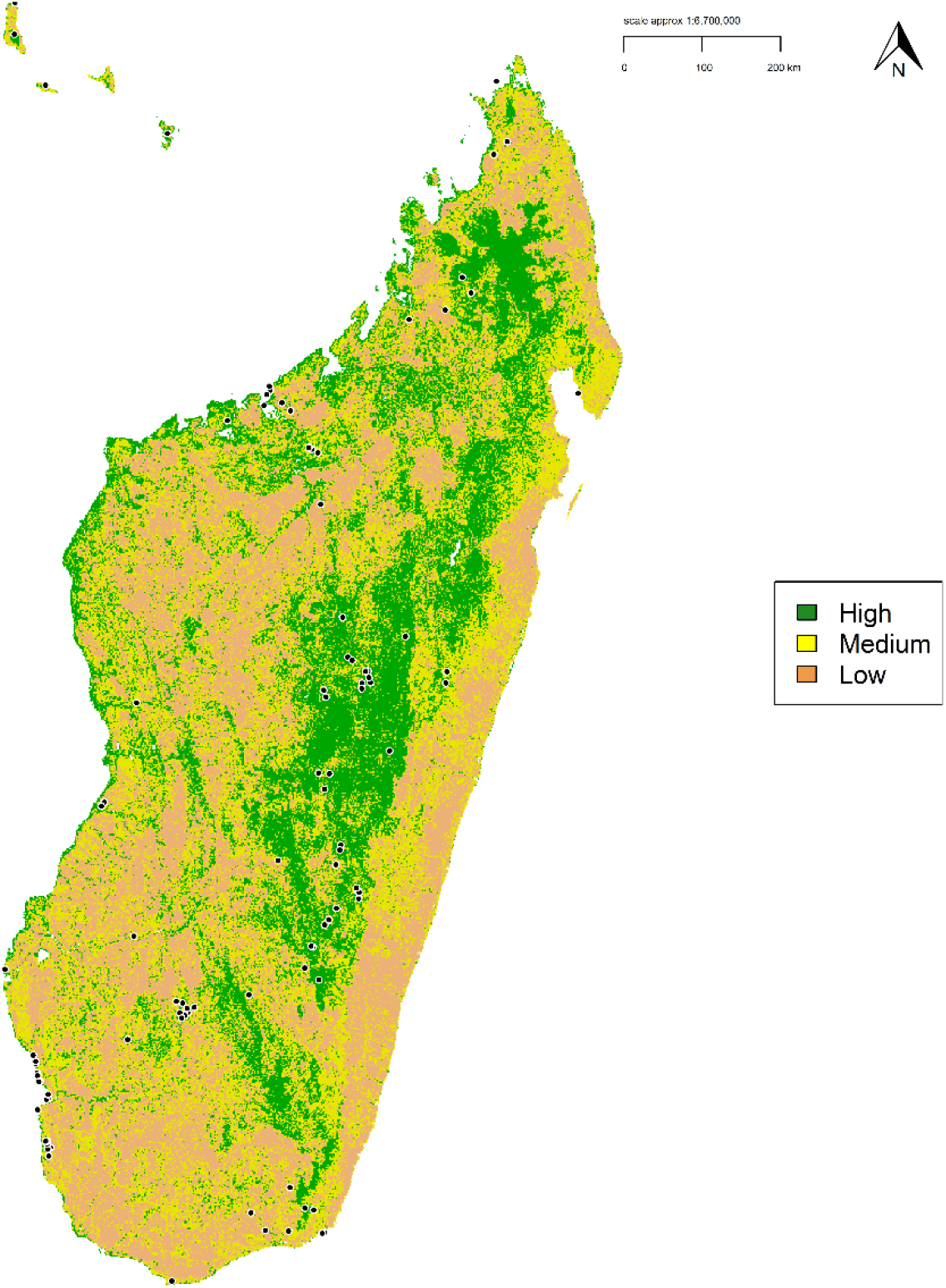
Reclassified discrete habitat suitability model for the Madagascar Peregrine Falcon. Map denotes the continuous prediction reclassifed into three discrete quantile threshold classes. Black points define Madagascar Peregrine occurrences.

## Discussion

Identifying species ecological requirements is fundamental to understanding species range limits and thus setting conservation planning priorities (Elith & Leathwick 2009; Lawler *et al*. 2011). Here, we provide the first detailed distribution map for the Madagascar Peregrine and establish this sub-species’ distributional constraints across its entire range using climatic, topographical, and landcover variables. Our results show predictive models can identify those areas of highest habitat suitability and infer the processes driving distribution for this sub-species in Madagascar. The core range for the Madagascar Peregrine extends across the central plateau of Madagascar, with a patchier distribution around coastal and lowland areas. The Madagascar Peregrine is more likely to be associated with high altitude, arid areas with high vegetation heterogeneity and herbaceous landcover but less likely in areas of cultivated land and mosaic forest. Based on our prediction of high-class habitat, we estimate there could be up to 300 pairs resident throughout their range, and thus we recommend the sub-species be classed as Vulnerable under IUCN guidelines.

Previous distribution maps for the Madagascar Peregrine have defined the entire land area across the sub-species range as its distributional limits (Ferguson-Lees & Christie 2005, Sinclair & Langrand 2013; BirdLife International 2019). White *et al*. (2013) reduced this to include a distributional area across the far northern and southern ends of Madagascar, and a separate area extending across central west Madagascar, with question marks in between these three areas. Our distribution estimates build on these previous attempts, providing both continuous and discrete predictions, and further detail than those previous range maps. We view these range maps as work in progress, which can be continuously updated as new occurrence data are collected. Indeed, using SDMs to map and update species range limits is gaining in use (e.g., Breiner *et al*. 2017; Ramesh *et al*. 2017; Da Silva *et al*. 2020). Using model-based interpolation from community science data offers a cost-effective and convenient method to assess and update the distribution status and range limits for this understudied sub-species, along with broad applications across other data-deficient taxa.

Few species occupy all areas with suitable resources and conditions, with areas of potential presence either occupied by closely related species, or unoccupied due to extirpation or failure to disperse (Anderson *et al*. 2002). For the Madagascar Peregrine, the largest area of high-class habitat extended across the central plateau, and further north to a smaller area of high-class habitat in the northern uplands in areas of high aridity. Increased precipitation is known as a strong negative predictor for Peregrine breeding success (Mearns & Newton 1988; Bradley 1997; Anctil *et al*. 2014) and may explain the positive relationship with areas of high aridity. Across their range the Madagascar Peregrine prefers areas of high vegetation heterogeneity, which may be a useful proxy associated with increased prey densities in highly heterogenous areas (Radeloff *et al*. 2019). Indeed, Peregrine population densities in tropical areas may be restricted by prey resource availability driven by habitat heterogeneity compared to temperate regions (Jenkins & Hockey 2001).

Though the Madagascar Peregrine would likely have evolved over the largely forested island of Madagascar prior to human-driven deforestation (White *et al*. 2013), elsewhere across their global range Peregrines are generally birds of open, treeless country, feeding largely on other avian species. Thus the strong association with areas of > 95 % herbaceous landcover in the form of open grasslands in Madagascar, albeit modified by humans, was expected. Interestingly, we found a negative coefficient relationship with cultivated land from the RSF, but a positive coefficient in the HSM with highest suitability in areas of 30 % cultivated land cover. We suspect that the use of a penalized GLM in the HSM prediction was able to identify this positive relationship by shrinking the other uninformative covariates, even though both models defined similar concave response curves.

Peregrine pair densities vary widely across their global range dependent on conditions and resources (Ratcliffe 1993; White *et al*. 2013). Our population size estimate ranging between 150-300 pairs based on inferred high-class habitat is similar to a previous estimate for the Madagascar Peregrine which used the entire terrestrial area of their range but with a density of one pair per 2000-4000 km^2^ (White *et al*. 2013). Estimating population density from SDMs is reliant on having sufficient and accurate occurrence data and selecting the most relevant predictors based on species biology (Franklin 2010; Oliver *et al*. 2012). Our method here is fairly coarse but gives a first estimate and a baseline from which to develop an integrated approach combining SDMs with population demography. For future range assessments, demographic response studies combined with Habitat Suitability Models are a promising method to further evaluate the relationship between population dynamics and habitat suitability (Bocedi *et al*. 2014). Further, modelling future distribution with both predicted prey distributions and land cover change may also yield useful insights into the future conservation status for the Madagascar Peregrine.

We recognise our results are reliant on the selected environmental covariates dependent on our definitions - animals may not experience the landscape as humans perceive it. Though we used a range of relevant abiotic and biotic predictors, including predictors derived from recent satellite remote sensing such as Dynamic Habitat Indices (Hobi *et al*. 2017, Radeloff *et al*. 2019), would further build on the predictive ability of the SDMs presented here. Spatial models have much scope for improvement (Engler *et al*. 2017; Fourcade *et al*. 2017; Guisan *et al*. 2017), but ultimately may also be limited by the amount and quality of occurrence data required (Beck *et al*. 2014; Neate-Clegg *et al*. 2020). Integrating multiple biodiversity data sources would seem a logical evolution for SDMs, particularly within a point process modelling (PPM) framework (Isaac *et al*. 2019). As shown here, the PPM framework using logistic regression produces reliable and useful predictions for species range limits in the absence of detailed distributional information for a little-studied raptor sub-species.

Improving our understanding of Peregrine habitat use is especially relevant for conservation because it may point towards potential population change and direct habitat management and protection. Developing a collaborative working relationship with field researchers and land managers feeding back occurrence data would further improve habitat use assessments. In particular, locations of known nest sites would help identify the specific environmental breeding requirements for the Madagascar Peregrine, as the occurrence dataset used here is mainly comprised of sightings. Given the discontinuous range of the Madagascar Peregrine, any breeding habitat studies would be more effective at local, landscape scales. Moreover, developing a participatory modelling process (Ferraz *et al*. 2020), where researchers, planners, and decision makers are all involved in the modelling process would be a significant step forward for raptor conservation in Madagascar.

Species ranges have an invariably discontinuous spatial arrangement, and this most likely applies to all taxa if mapped at fine scales (Ladle & Whittaker 2011). In the absence of human disturbance, discontinuous distributions are a consequence of patchy environments, so it follows that species ranges will follow the latter (Riddle *et al*. 2011). Range occupancy is also scale dependent, with varying scales of spatial pattern in individual species (Levin 1992; Garshelis 2000). Refining broad scale modelling to localized regional studies would be a logical next step. Exploratory ground-truthing surveys to validate the models would also be of benefit to conservation managers and modellers alike. Many developing tropical countries lack extensive resources for biodiversity management but are generally the regions of the planet with highest biodiversity. Vast wilderness areas make systematic sampling for many species almost impossible, and that is where spatial models are required to fill knowledge gaps and determine potential distributional areas as demonstrated here.

## Acknowledgements

We thank all individuals and organisations who contributed occurrence data to the Global Raptor Impact Network (GRIN) information system. We thank the M.J. Murdock Charitable Trust for funding and the Information Technology staff and Research Library interns at The Peregrine Fund for support.

## Data Accessibility Statement

Upon acceptance the data that support this study will be made openly available on the data repository *figshare*

## Supplementary Material

### Appendix 1

#### Supplementary Tables

**Table S1.** Multi-collinearity test using stepwise elimination Variance Inflation Factor (VIF) analysis. Variables with VIF < 5 have low correlation with other variables, and thus are suitable for inclusion in calibration models when further evaluated for ecological relevance.

| Covariate | R<sup>2</sup> | Tolerance | VIF |
| --- | --- | --- | --- |
| Aridity Index | 0.308 | 0.692 | 1.444 |
| Elevation | 0.244 | 0.756 | 1.323 |
| Herbaceous | 0.227 | 0.773 | 1.293 |
| Heterogeneity | 0.224 | 0.776 | 1.289 |
| Mosaic forest | 0.180 | 0.820 | 1.220 |
| Cultivated | 0.159 | 0.841 | 1.189 |

**Table S2.**
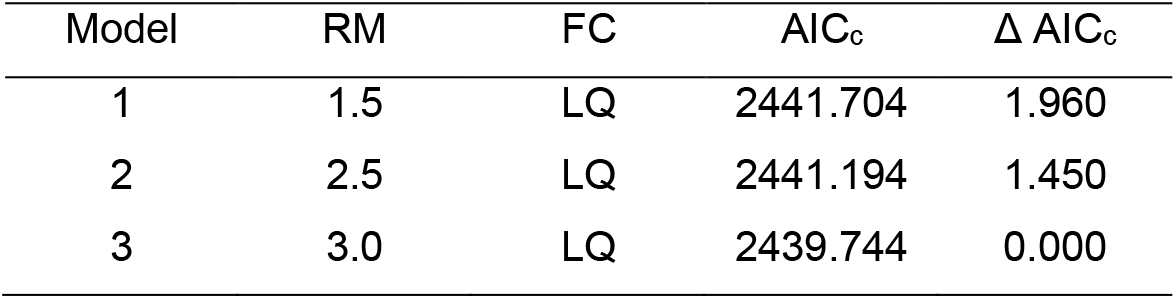
Model selection metrics for all candidate models with ΔAIC_c_ < 2. RM = regularization multiplier (β), FC = feature classes, LQ = Linear, Quadratic.

